# An extension of Wang’s protein design model using Blosum62 substitution matrix

**DOI:** 10.1101/2021.06.07.447415

**Authors:** Amin Rahmani, Fatemeh Zare Mirakabad

**Affiliations:** Amirkabir University of Thechnology

## Abstract

Humans life depends on the functionality of molecules in the body. One of these essential molecules is the protein that plays a vital role in our life, such that its malfunction can cause severe damages. Such roles make protein structure and its functionality necessary to understand. One of the problems that help us understand the relation between protein structure is the well-known protein design problem which attempts to find an amino acid sequence that can fold into a desired tertiary structure. However, despite having an acceptable accuracy in protein design, this accuracy is an identical percentage of amino acid retrieving. At the same time, it is well-known that amino acids can replace each other in evolution while the function and structure of protein stay the same. Thus the designed sequence does not have the opportunity to be close to the target in the evolutionary aspect. This paper presents an extension to Wang’s deep learning model, which uses evolutionary information in the Blosum62 substitution matrix to take amino acid replacement probability into account while designing a sequence.

## Introduction

Proteins are one of the most important molecules in life. Different functionalities, e.g., helping the olfactory system in sensing smells[1], catalyzing metabolism reactions[2], and a vast majority of other significant roles in the human body, made this macromolecule an essential topic of study in biology. The functionality of protein comes from its most significant structure, the tertiary structure[3]. This structure defines an almost unique shape for the protein. This unique shape determines the ligands that the protein can interact with and how robust their binding is. Thus, misfolding in a protein causes issues with binding that affect its functionality. In addition, some neurological diseases arise from protein misfolding, e.g., Alzheimer’s, Parkinson’s, and Huntington’s disease[4]. With all this importance in mind and PDB saturation in tertiary structures, understanding the relationship between primary and tertiary structures helps us in protein tertiary structure prediction and genome sequence functionality prediction. One of the approaches for understanding the relationship between primary and tertiary structures is discovering a sequence of amino acids that can get a desired tertiary structure, referred to as the protein design problem (PDP). Some applications of PDP are designing proteins that can interact with specific targets[5] and designing biosensors[6].

The PDP takes a backbone-only tertiary structure of a protein as input and produces a sequence of amino acids as its output. The objective of the problem is that when the output amino acid sequence folds into a 3-Dimensional shape, the folded structure be similar to the input structure. PDP is an NP-Hard problem[7]. For a more solid perception let’s assume a polypeptide chain with 100 residues, in this case, there are 100^20^ unique possible amino acid sequences for this chain, and thus discovering the most compatible with desired structure is hard to solve.

There are three broad classes of PDP algorithms, exact algorithms, heuristic algorithms, and machine learning. The first class contains exact algorithms such as dead-end elimination that can provide a guaranty to find a solution if it exists, but there is no guaranty for runtime in this class of algorithms; therefore, running such algorithms for PDP costs high in terms of runtime[8]. The second class of algorithms uses a heuristic algorithm to optimize a designed energy function and find a sequence that minimizes this energy function. PDP models abundantly use heuristic algorithms like Monte Carlo; however, running this kind of optimization algorithm needs a vast iteration at each design, and also, besides this runtime issue for heuristic algorithms, it is hard to design energy functions[9]. For example, PDP tools like RosettaDesign[10] and EvoDesign[11]use this approach. The last class of algorithms uses machine learning methods, especially Deep Learning methods. Deep learning, just like other machine learning methods, uses data to learn a mapping between the input and desired output. Although these algorithms need a vast amount of data and considerable time for training concerning network architecture, they are more desirable because of their speed in usage.

Machine learning approaches for PDP use previously collected protein structures and sequences to learn protein design. Thus, there is no need for designing an energy function. Due to machine learning capability, there were efforts, during recent years, to overcome the protein design problem using this technique and made considerable progress. In 2014 Li et al. used a simple neural network with two hidden layers and extracted structural features as input[12]. After that, in 2018, O’Connell et al. developed a deep neural network with higher accuracy and more extracted features from data for the network input[13]. In the same year, Wang et al. provided a more complex model and the same method for input extraction from data[14]. The last developed model called DenseCPD by Qi et al. in 2020 uses a residual convolutional network and a representation of 3D structure to predict corresponding amino acid type for each residue[15]. These models, despite having acceptable accuracies lack quality in produced sequences.This inadequateness in the quality of generated sequences has two main reasons. The first is a supervised training method where the targets for output are one-hot encodings of amino acid classes and the second reason is lack of information in the input data.

One-hot encoded targets for training cause the network to restore the wild-type sequence of structure, although some non-identical protein sequences have similar foldings due to their mutations in evolution. Also, feature selection is difficult to do, and selecting these features as a representation of a residue of the protein structure may not be a valuable input. The first three models have feature selection and identical retrieval of amino acids, and the last model, Dense CPD, despite having a new representation of protein structure that seems informative yet tends to retrieve identical amino acids to the target training sequence.

The fact that each amino acid tends to replace specific amino acids can be interpreted by studying the evolution of protein sequences. This information is available in substitution matrices. Substitution matrices like Blosum encompass helpful information from protein sequence evolutionary data, which can help us understand which amino acids have the desire to take place at another amino acid position in a sequence. Substitution matrices take amino acid substitution scores and conserver blocks of protein into account and generate a matrice with substitution scores for each two amino acids based on evolutionary data. Therefore, amino acids with the desire to stay conserved have a lower substitution score[16].

In this paper, we review Wang’s model in detail and then extend this model to obtain generated sequences with higher quality.

## Material and Methods

As illustrated in Figure 1 Protein Design Diagram, we can explain PDP as a problem with inputs, outputs, and goals. The input is a backbone-only model of tertiary structure for a protein, and the output is a sequence of amino acids. The aim is to design the output sequence so that the resulting structure is similar to the input structure when it folds into a tertiary structure.

**Figure 1.**
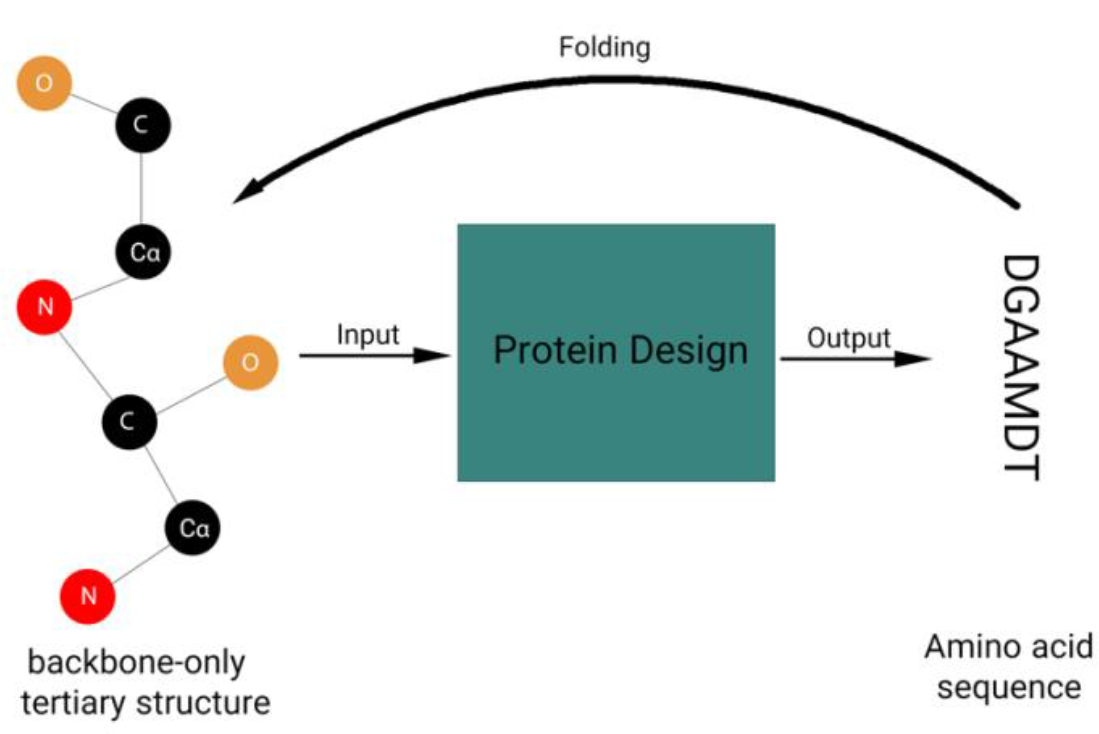
Protein Design Diagram

Among the provided machine learning methods for PDP so far, Wang’s model has the second-highest accuracy among reviewed models after DenseCPD, therefore due to the lack of resources to train DenseCPD, we decided to extend Wang’s model.

In the following sections, first, we discuss data collection, and after that, we review Wang’s model. After Wang’s model, we present an extension to Wangs model

### Dataset

Data collection happens by retrieving protein structure and corresponding sequences from PDB with the following criteria as Wang’s model does. (1) The structure determination method is x-ray crystallography, (2) The resolution of the tertiary structure is better than 2Å, (3) The length of the protein sequence is more than 50 amino acids, (4) The entry does not contain any DNA or RNA molecules, and (5)Amino acid sequences of all found entries have less than 30% pairwise identity.

Furthermore, by cross-referencing the retrieved data with the OPM database[17], membrane proteins can be found and removed. Later, each entry containing D-amino acids vanishes from the dataset, and the non-amino acid residues of each protein also exclude from the structure. In the next step, each protein with a sequence length of *L*_*s*_ split into *L*_*s*_ clusters, where each corresponds to one of the residues. Each cluster contains a target residue and its 15 nearest neighbors regarding the *c*_*α*_ − *c*_*α*_ distance. For each cluster, all the neighbor and target residues rotate and translate such that the *c*_*α*_ atom of the target residue locates on the (0,0,0) point, the *N* atom of the target residue lies on the −*x* axis, and the *C* atom of the target residue takes place in the *z* = 0 plane.

### Wang’s model

In this section, we review the input, output, and architecture of Wang’s model in detail.

#### Input

For input, feature extraction happens on each cluster, and each one of the clusters would have two types of feature sets; one set of features for the target residue of that cluster and the other set of features correspond to each of the neighbors in the cluster. Target residue feature set contains sine and cosine of three backbone dihedral angles *ϕ, ψ*, and *ω*, the total solvent accessible surface area(SASA) of backbone atoms, and the three stated secondary structures (helix, sheet, or coil) represented with a one-hot vector. As for the neighboring residues, the feature set for each one contains sine and cosine of three backbone dihedral angles phi, psi, and omega, the total solvent accessible surface area of backbone atoms, three stated secondary structures (helix, sheet, or coil) represented with a one-hot vector, *C* − *C* Euclidean distance to target residue, unit *C* − *C* vector from the target residue to the neighboring residue, unit *C* − *N* vector in the under process neighbor residue, unit *C* − *C* vector in the under process neighbor residue, and number of hydrogen bonds between the target and the neighbor residue. Thus, the feature extraction procedure for each cluster results in 10 features for the target residue and 24 features for each neighboring residues.

#### Output

We perform one-hot encoding of the amino acid type for each cluster’s target residue as output targets. Thus, the model outputs a vector of size 20 that we interpret as probabilities of different amino acid types for the input cluster.

#### Model

The presented model by wang et al., as presented in Figure 2: Architecture of Wang’s model has two sub-networks and three final layers before the output layer. The sub-networks are called residue probability network and weight network. The residue probability network tends to find primal probabilities for the class of target residue by seeing this residue and one of its neighbor residues features, and the Weight network produces a weight by considering the same input as the residue probability network as well. The output of two subnetworks is then multiplied by each other and kept as part of the input for later layers. This procedure executes concurrently for 15 nearest neighbors of the target amino acid, and then the multiplied result of all is concatenated to each other. The concatenated result feeds into three layers of densely connected layers, and at the end, a softmax layer containing 20 nodes outputs a probability vector.

**Figure 2:**
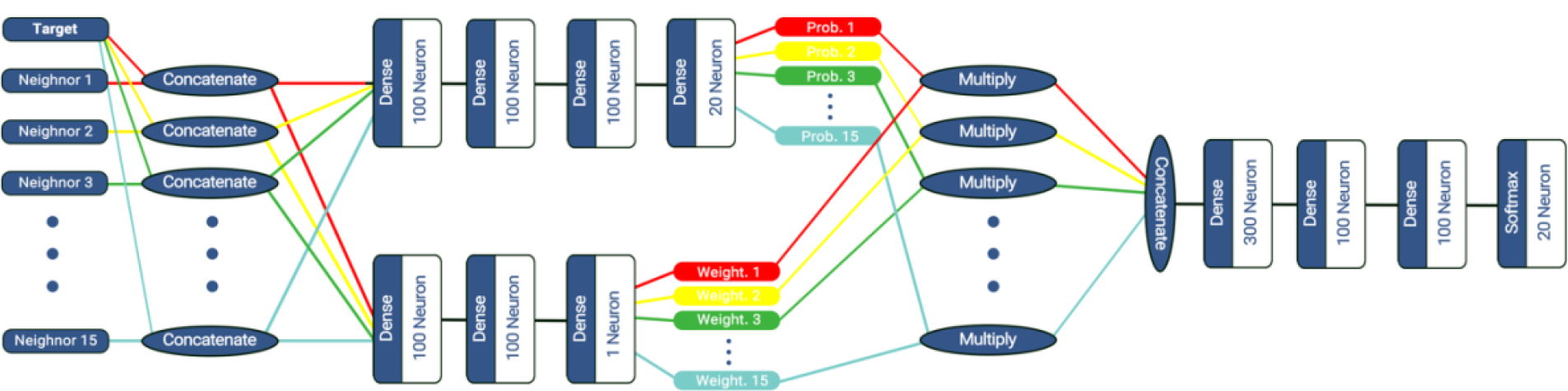
Architecture of Wang’s model

This network mimics the behavior of convolutional networks, which sees not only the target input but also its neighbors.

### Extending wang’s model

#### Input and architecture

The input and architecture for our extended model is the same as Wang’s model and no change happens.

#### output

A solution to the quality of sequences lies in the way they evolve. So instead of training the network with one-hot encoded targets, it is rational to use a vector that considers other probabilities. We use the characteristic of the Blosum matrices and present a new target of training that contains probabilities of multiple amino acid classes. We chose Blosum62, which contains substitution information from proteins with less than 62% identity[18].

To present these scores as targets of training the network, preprocessing is necessary. To transform scores into probabilities, we applied the softmax function to each row of the Blosum62 matrix. Eventually, these converted rows are considered as targets and replaced with one-hot encoded vectors for loss calculation.

## Results

We used Keras^3^ for the implementation of both of the networks, as the Wang et al. paper suggested. Also, all activations are ReLU, the used loss is categorical cross-entropy with a learning rate of 0.01 and a Nesterov momentum of 0.9, and the batch size is 128.

Because the number of clusters was numerous, 100,000 clusters were selected randomly and split into three non-redundant train, test, and validation datasets with sizes of 70,000, 15,000, and 15,000. The same results as the original model were first regenerated. Then, after modifying and training the network on the same data, we compared the results of both the original and extended models on the test set.

**Table 1:** Acuuracy of models on amino acids identical prediction

|  | Test Accuracy |
| --- | --- |
| Wang's model | 36.39% |
| Extended model | 44.89% |
| DenseCPD | **55.53%** |

Comparision shows a remarkable improvement in sequence identity and accuracy compared to Wang’s model and acceptable accuracy regarding the data sample and training time compared to the DenseCPD.

Furthermore, we design and analyze sequences for two proteins that do not exist in the train, test, and validation sets. These two proteins have PDB ID **1HOE** and **1QV1** with lengths of 74 and 187 respectively.

Analyzing includes different tools and comparison:

1. aligning the designed and natural protein sequences using PSI-Blast
2. predicting the 3-Dimensional structures, and aligning them with the original structure using I-Tasser and TM-Align
3. comparison of extracted secondary structures, dihedral angles, and SASA from predicted tertiary structures using DSSP
4. predicting the subcellular localization using LocTree3
5. examining predicted binding site of each sequence using ProNA2020

### PSI-Blast alignment

PSI-Blast can align multiple sequences and returns alignment score, E-value, and identity of sequences[19]. The alignment score is the total score of alignment calculated by a substitution matrix, and the E-value reflects the probability of finding a match for the query sequence in the database. As we can see in Table 2: Analyzing sequences quality using PSI-Blast alignment with natural sequence, there is a significant improvement in both score and E-value for the sequence designed by the extended model. Even though the identity of Wang’s generated sequence is higher than the extended model for 1QV1, we can see the extended model gets higher scores and lower E-value. For the other protein, 1HOE, Wang’s designed sequence has a very shallow alignment; therefore, no score and E-value are available, and also, the identity is lower than the extended model’s designed sequence.

**Table 2:** Analyzing sequences quality using PSI-Blast alignment with natural sequence

|  | 1QV1 |  |  | 1HOE |  |  |
| --- | --- | --- | --- | --- | --- | --- |
| Model | Score | E-Value | Identity | Score | E-Value | Identity |
| Wang | 31.2 | $1 \times e^{-5}$ | **52.50%** | - | - | 32.34% |
| Extended | **164** | **$4 \times e^{-56}$** | 49.09% | **65.5** | **$7 \times e^{-21}$** | **50%** |

### 3-Dimensional structures comparison

Since protein folding is also an NP-hard problem and there is no deterministic solution, we use a folding prediction tool named I-Tasser[20]. For comparison, first, we use the I-Tasser to predict a structure for the natural sequence of the protein, and then we predict the structure for the two designed sequences. In all predictions, we use the first model that I-Tasser provides. Next, we use TM-Align[21], a tool for tertiary structure alignment, and align predicted tertiary structures for natural and the two designed sequences to measure the closeness of predicted 3-Dimensional structures. Two factors describe this similarity; root-mean-squared deviation (RMSD) and alignment score (TM-Score). A lower RMSD indicates a better match of atomic positions, and a higher TM-Score means the two aligned structures are close in terms of quality.

**Table 3:** Comparison of predicted structures with the natural structure I-Tasser and TM-Align

| Sequence | 1QV1 |  | 1HOE |  |
| --- | --- | --- | --- | --- |
|  | RMSD | TM-Score | RMSD | TM-Score |
| Natural | 0.71 | 0.96057 | 0.49 | 0.97615 |
| Rosetta | **0.64** | **0.98886** | 2.14 | 0.74829 |
| Extended | 0.74 | 0.74829 | **0.77** | **0.96262** |

### Secondary structures, dihedral angles, and SASA comparison

Next, we extract dihedral angles phi and psi, solvent accessible surface area, and secondary structures from predicted structures using the DSSP program[22]. The dissimilarity function used to compare angles and SASA is root-mean-squared-error (RMSE) because the numbers are continuous; consequently, a lower RMSE presents a higher similarity.

**Table 4:** RMSD of extracted dihedral angles and SASA to the dihedral angles and SASA of real structure using DSSP

| Model | 1QV1 |  |  | 1HOE |  |  |
| --- | --- | --- | --- | --- | --- | --- |
| | $RMSE_{SASA}$ | $RMSE_{\phi}$ | $RMSE_{\psi}$ | $RMSE_{SASA}$ | $RMSE_{\phi}$ | $RMSE_{\psi}$ |
| Natural | 42.62 | 36.69 | 40.12 | 41.40 | 56.22 | 84.64 |
| Rosetta | **40.19** | 48.59 | **49.18** | 45.05 | 64.38 | 105.2 |
| Extended | 42.77 | **41.61** | 60.10 | **43.92** | **55.63** | **96.70** |

Continuing the comparisons, next is the secondary structure. For this comparison, first, we use the DSSP tool and extract secondary structures of retrieved PDB entry for protein, predicted structure for the retrieved PDB, and the predicted structure of the designed sequences. For each structure, we build a sequence of its residues secondary structure and then measure the accuracy. Besides accuracy, we also measure precision and recall to show the reliability of the accuracy.

**Table 5:** Accuracy, precision, and recall for the extracted secondary structures from predicted structures and natural secondary structures

| Model | 1QV1 |  |  | 1HOE |  |  |
| --- | --- | --- | --- | --- | --- | --- |
|  | Accuracy | Precision | Recall | Accuracy | Precision | Recall |
| Natural | 86.63% | 81.46% | 86.48% | 67.56% | 72.86% | 72.04% |
| Rosetta | 87.70% | 79.63% | 86.23% | 58.10% | 52.82% | 54.67% |
| Extended | **88.23%** | **85.96%** | **90.93%** | **70.27%** | **75.76%** | **68.50%** |

### Subcellular localization prediction

For the subcellular localization prediction, we use a tool named LocTree3, which predicts the subcellular localization of input protein sequence among 18 classes for eukaryotes[23]. We report confidence alongside the prediction of the GO terms.

**Table 6:**
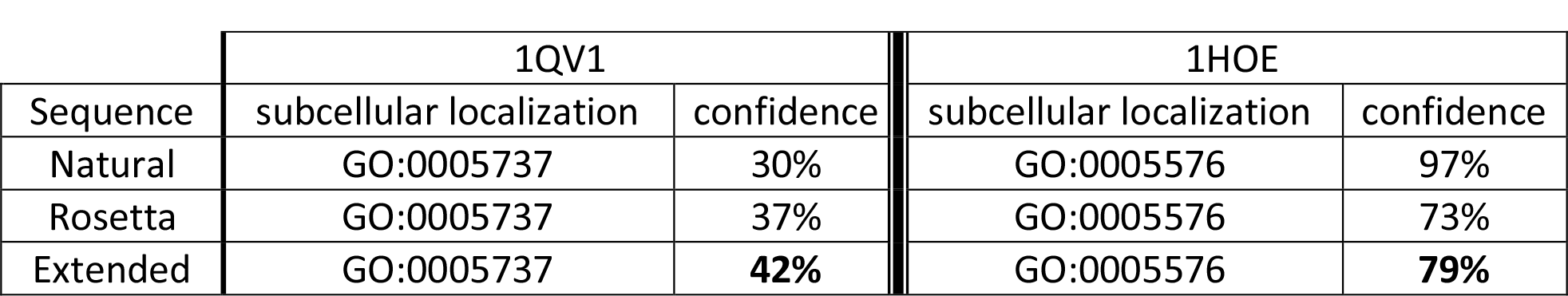
Subcellular localization prediction for natural sequence and designed sequences

### Binding sites prediction and comparison

Eventually, we compare the binding sites of each sequence. To do so, first, we use ProNA2020 to predict whether each position of the sequence does belong to a binding site or not and generate a binary binding site sequence[24]. Afterward, we compare the binding site sequence of designed sequences with the binding site sequence of the natural protein sequence. As a measurement, we use accuracy, which shows us how good the predicted binding sites of the designed protein can be close to the prediction of sites for the natural sequence.

**Table 7:** Accuracy of predicted binding sites comparing to predicted binding site of natural sequence

|  | 1QV1 | 1HOE |
| --- | --- | --- |
| Rosetta | 91.89% | - |
| Extended | **93.24%** | **68.91%** |

As we can see for most of the comparisons, our extended model generates a well-quality sequence and can do better than the Rosetta. However, Rosseta has a better performance on the 1QV1, which is almost twice longer than 1HOE. Nevertheless, the extended deep learning model has the advantage of run time, which is significantly lower than the Rosetta and generates sequences that are slightly different from Rosetta generated sequences in terms of quality.

## Conclusion

In this paper, we investigated the protein design problem and its methods. One of the methods used machine learning, but despite having more than 30% identity of output to the natural sequence of the tertiary input structure, the generated sequences had low quality such that no natural protein sequence would have those characteristics. We provided an extension to this model, named Wang’s model, and with such a minimal extend achieved much better results.

## Footnotes

3 https://keras.io/

## Notes

### Competing Interest Statement

The authors have declared no competing interest.

